# Comparative Transcriptomes Analysis of *Taenia pisiformis* at Different Development Stages

**DOI:** 10.1101/490276

**Authors:** Lin Chen, Jing Yu, Jing Xu, Wei Wang, Lili Ji, Chengzhong Yang, Hua Yu

## Abstract

To understand the characteristics of the transcriptional group of *Taenia pisiformis* at different developmental stages, and to lay the foundation for the screening of vaccine antigens and drug target genes, the transcriptomes of adult and larva of *T. pisiformis* were assembled and analyzed using bioinformatic tools. A total of 36,951 unigenes with a mean length of 950bp were formed, among which 12,665, 8,188, 7,577, and 6,293 unigenes have been annotated respectively by sequence similarity analysis with four databases (NR, Swiss-Prot, KOG, and KEGG). It should be noted there are 5,662 unigenes that share good similarity with the four databases and get a relatively perfect functional annotation. Besides, a total of 10,247 differentially expressed genes were screened. To be specific, 6,910 unigenes were up-regulated in the larva stage while 3,337 were down-regulated in the adult stage. To sum up, this study sequenced and analyzed the transcriptomes of the larval and adult stages of *T. pisiformis*. The results of differentially expressed genes in these two stages could provide basis for functional genomics, immunology and gene expression profiles of *T. pisiformis*.

## 1. Introduction

Cysticercosis is one of the common parasitic diseases in rabbits, caused by the metacestode of *Taenia pisiformis* (Cestoidea; Cyclophyllidea; Taeniidae), also known as *Cysticercus pisiformis*. The adult *T. pisiformis* often parasitized the small intestine of canines and felines, such as dogs, wolves, jackals, foxes, raccoon dogs and cats (Owiny, 2001; Foronda and Valladares et al., 2003; Saeed and Maddox et al., 2006; Martinez and Hernandez et al., 2007; Lahmar and Sarciron et al., 2008; Bagrade and Kirjusina et al., 2009; Jia and Yan et al., 2010). The larvae (Cysticercus pisiformis) often inflict the liver capsule, greater omentum, mesentery and rectal serous membrane of rabbits, squirrels, mice, and other rodents (Zhou and Du et al., 2008). *T. pisiformis* can cause serious health problems and even death (Yang and Fu et al., 2012).

So far, there have been a lot of research on *T. pisiformis*, including its biological characteristics (Chen and Yang et al., 2015), population genetic diversity (Yang and Ren et al., 2013), gene function (Yang and Chen et al., 2013; Chen and Yang et al., 2014; 2014; Chen and Yang et al., 2016) and preventative measures against it (Chen and Yang et al., 2014). However, although the transcriptome of the adult of *T. pisiformis* has been determined before (Yang and Fu et al., 2012), there is no report on comparative transcriptomics of *T. pisiformis* at different developmental stages. Therefore, the exploration of gene expression patterns in different developmental stages of *T. pisiformis* will contribute to elucidating the mechanism of the infection of the host and provide an important basis for the prevention and control of these tapeworms. Yet, due to the limitation of materials and research methods, the infection mechanism of *T. pisiformis* is still unclear. In recent years, high throughput sequencing technology has been widely accepted thanks to its various advantages, such as large number of data, high accuracy, high sensitivity and low running cost. It has been used in many parasite species such as *Itch mite* (He and Xu et al., 2016), *Sarcoptes scabiei* (He and Gu et al., 2017), *Dirofilaria immitis* (Fu and Lan et al., 2012), *Taenia ovis* (Zheng, 2017), *Echiococcus graulosus* (Ju, 2013), *Taenia multiceps* (Li and Zhang et al., 2017) and others (Kolev and Franklin et al., 2010; Cantacessi and Young et al., 2011; Sorber and Dimon et al., 2011; 2012; Schicht and Qi et al., 2014). By using this method, it is possible to understand the molecular mechanism of specific biological processes and understand the overall level of gene expression at different stages of the parasite. In the current research, the Illumina sequencing techniques were applied to study the transcriptome of the two different developmental stages of *T. pisiformis*. By clarifying the gene expression status in the adult and larva of *T. pisiformis*, this research will lay a foundation for the diagnosis, prevention and treatment of Cysticercosis.

## 2. Results

### 2.1. Illumina sequencing and assembly

RNA-Seq generated approximately 20 to 26 million raw sequence reads for each of the six cDNA libraries obtained from *T. pisiformis* (three from adult stage and three from larva stage). The average GC content was 47.25% (Table 1). All of the clean reads were assembled into a transcriptome, which was used as a reference sequence for further analyses. In total, 36,951 unigenes were detected in all transcriptomes (Table 2). The mean length and N50 were 950bp and 1,998bp, respectively (Table 2). According to the statistics of length distribution, 13,759 unigenes (43.50%) ≥ 500bp, and 8,975 (24.29%) ≥ 1,000bp (Figure 1, Additional file 1: Table S1). The transcriptome raw reads dataset obtained has been submitted to the NCBI Short Read Archive (http://www.ncbi.nlm.nih.gov/Traces/sra_sub/sub.cgi) with the accession number: SUB4234089.

**Table 1.**
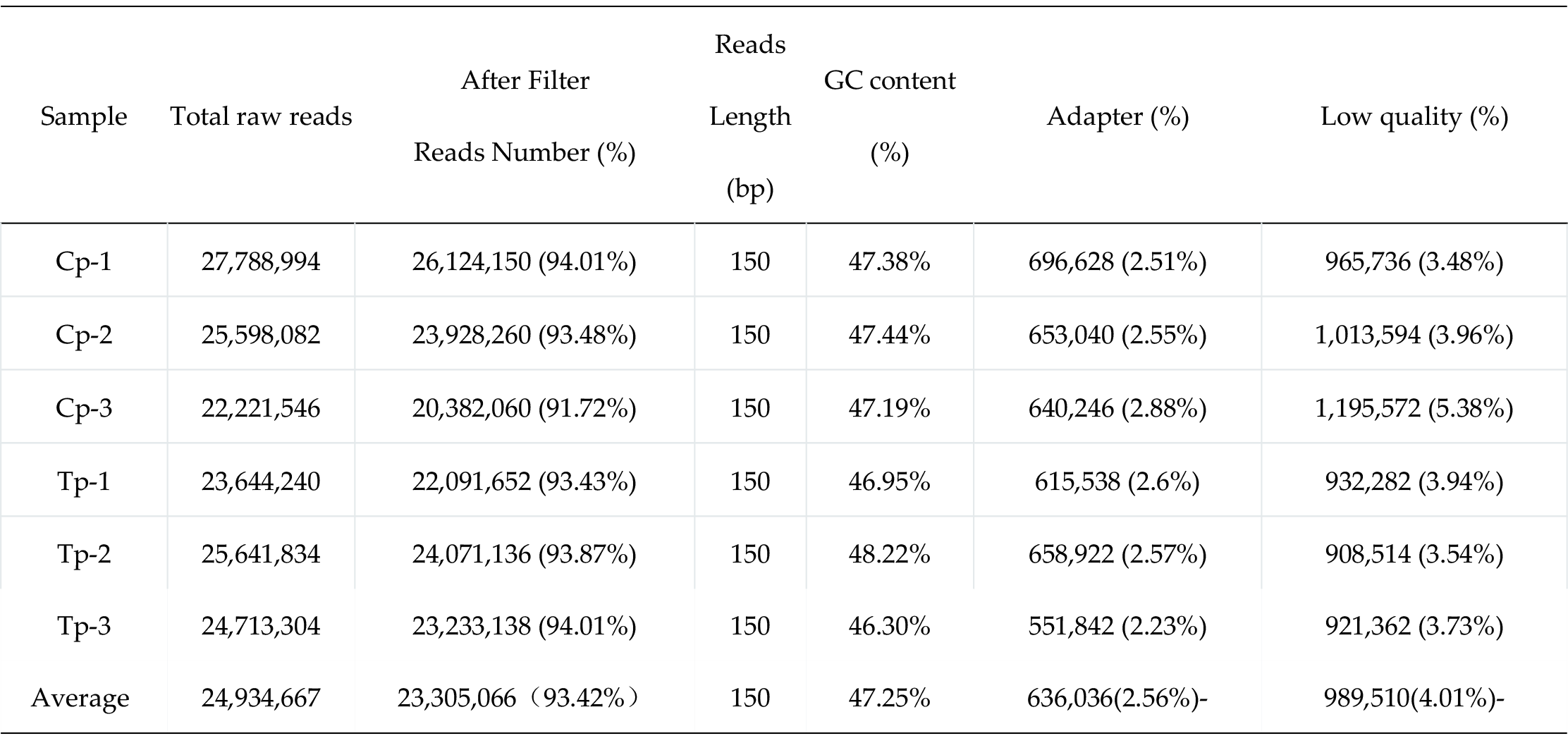
Summary of transcriptome data for adult (Tp) and larva (Cp) of *T. pisiformis*

**Table 2.** Filtered results of transcriptome data

| Genes<br>Number | GC<br>percentage<br>(%) | N50(bp) | Max length<br>(bp) | Min length<br>(bp) | Average<br>length (bp) | Total<br>assembled<br>bases |
| --- | --- | --- | --- | --- | --- | --- |
| 36,951 | 46.54 | 1,998 | 26,041 | 201 | 950 | 35,115,151 |

**Figure 1.**
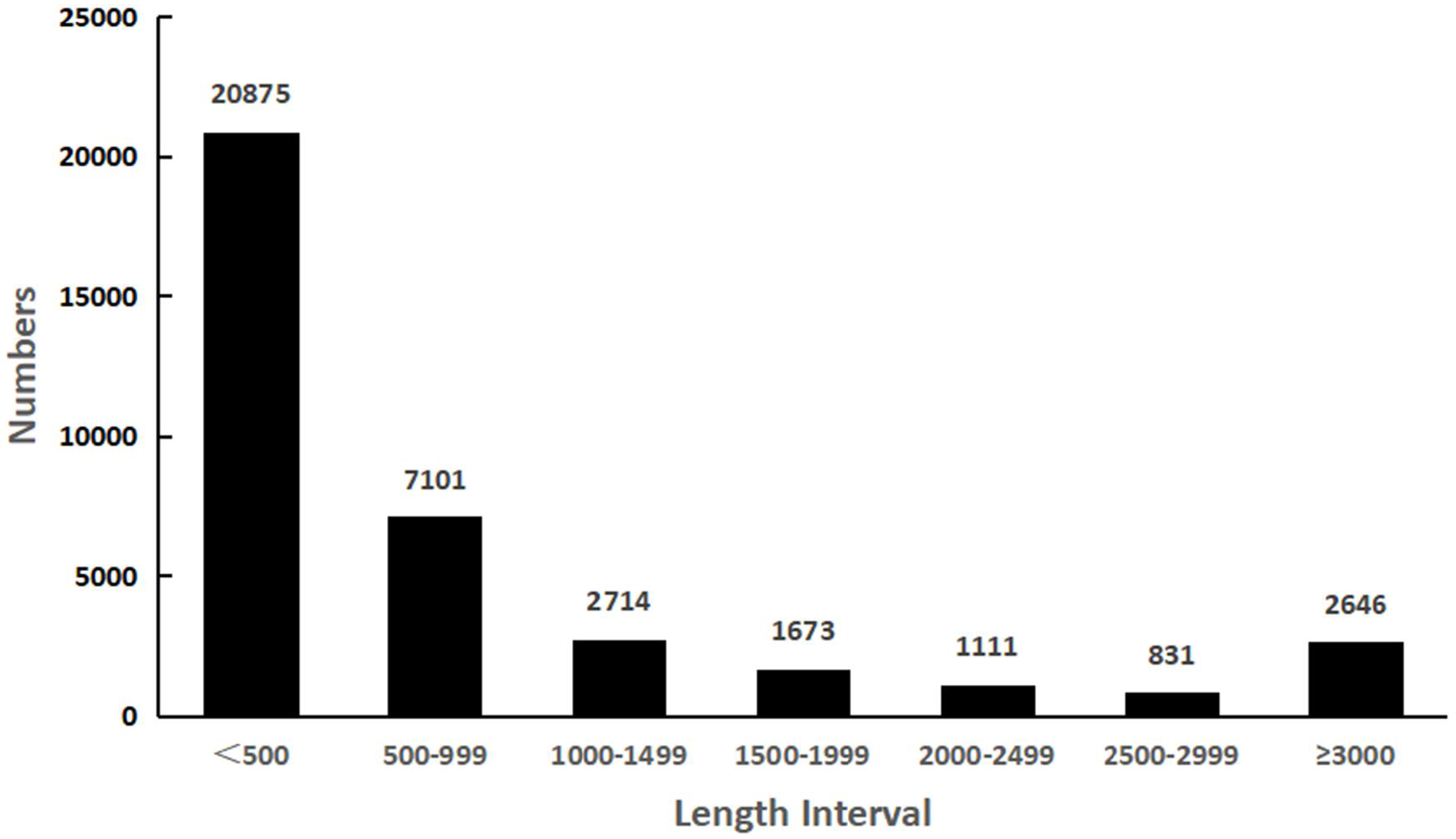
Length distribution of *T. pisiformis* unigenes

### 2.2 Functional annotation of unigenes

A total of 12,844 unigenes (34.76% of all unigenes) were annotated by the Nr (12,665, 34.28%), Swiss-Prot (8,188, 22.16%), KEGG (6,293, 17.03%) and KOG (7,577, 20.51%) protein databases using Blastx, and 5,662 unigenes were similar to the sequences of all these four databases (Figure 2). Rest of the unigenes (24,107) failed to match against any sequence with E-value < 10^− 5^. Homologous genes came from several species, with 59.01% of the unigenes having the highest homology to genes from *Echinococcus granulosus* (7,474, 59.01%), followed by *Echinococcus multilocularis* (3,654, 28.85%), *Hymenolepis microstoma* (557, 4.4%), *Daphnia magna* (72, 0.57%), *Taenia solium* (67, 0.53%) and other species (841, 6.64%).

**Figure 2.**
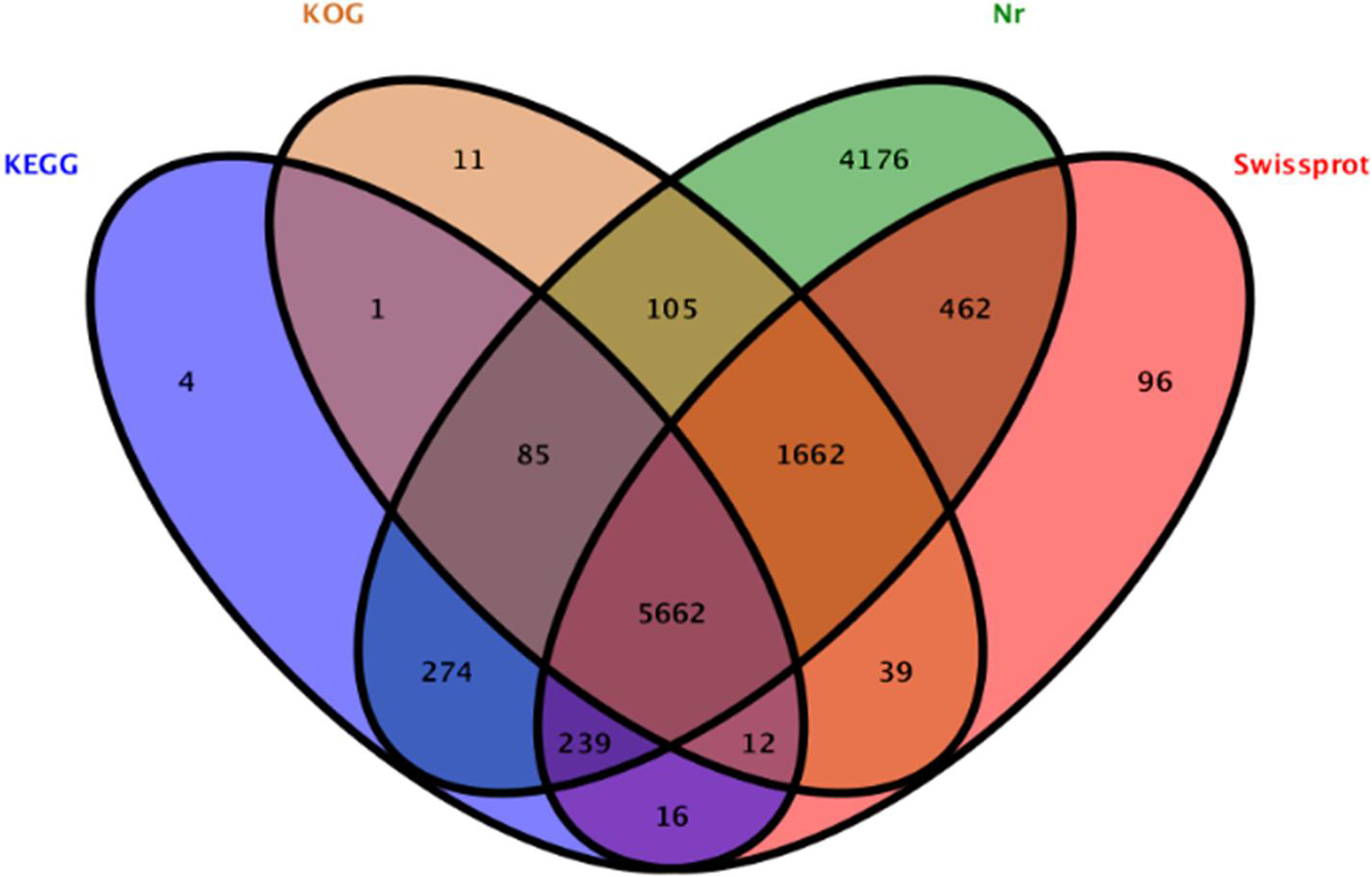
Wayne diagrams of four database annotations

The analysis of GO function annotation provided a functional classification and enrichment analysis for differentially expressed genes (DEGs) (Li and Zhang et al., 2017). As shown in Figure 3, 16,269 unigenes can be classified into 25 independent KOG functional items. “Signal transduction mechanisms” is one of the largest clusters, including 2,882 (17.7%) unigenes. The second and third largest ones are “general function prediction only” (containing 2,393 unigenes, 14.7%) and “modification after translation, transformation” (containing 1699 unigenes, 10.4%). “Cell motility” had the lowest number of related genes (only 34). In the transcriptome, the items that involve the growth of *T. pisiformis* include: “carbohydrates (2.24%)”, “transport and metabolism of amino acid transport and metabolism (1.46%)”, “nucleic acid transport and metabolism (1.03%)”, “lipid transport and metabolism (1.82%)”, “other material metabolism and signal transduction mechanism (17.71%)”, “transcription (6.62%)”, “translation, the ribosome structure and biological transformation (4.88%)”, “protein modification after translation, protein conversion, molecular partner (10.44%)”, “coenzyme transport and metabolism (0.48%)”, “secondary metabolites biosynthesis, transportation and catabolism (0.47%)” and “defense mechanism (0.54%)”.

**Figure 3.**
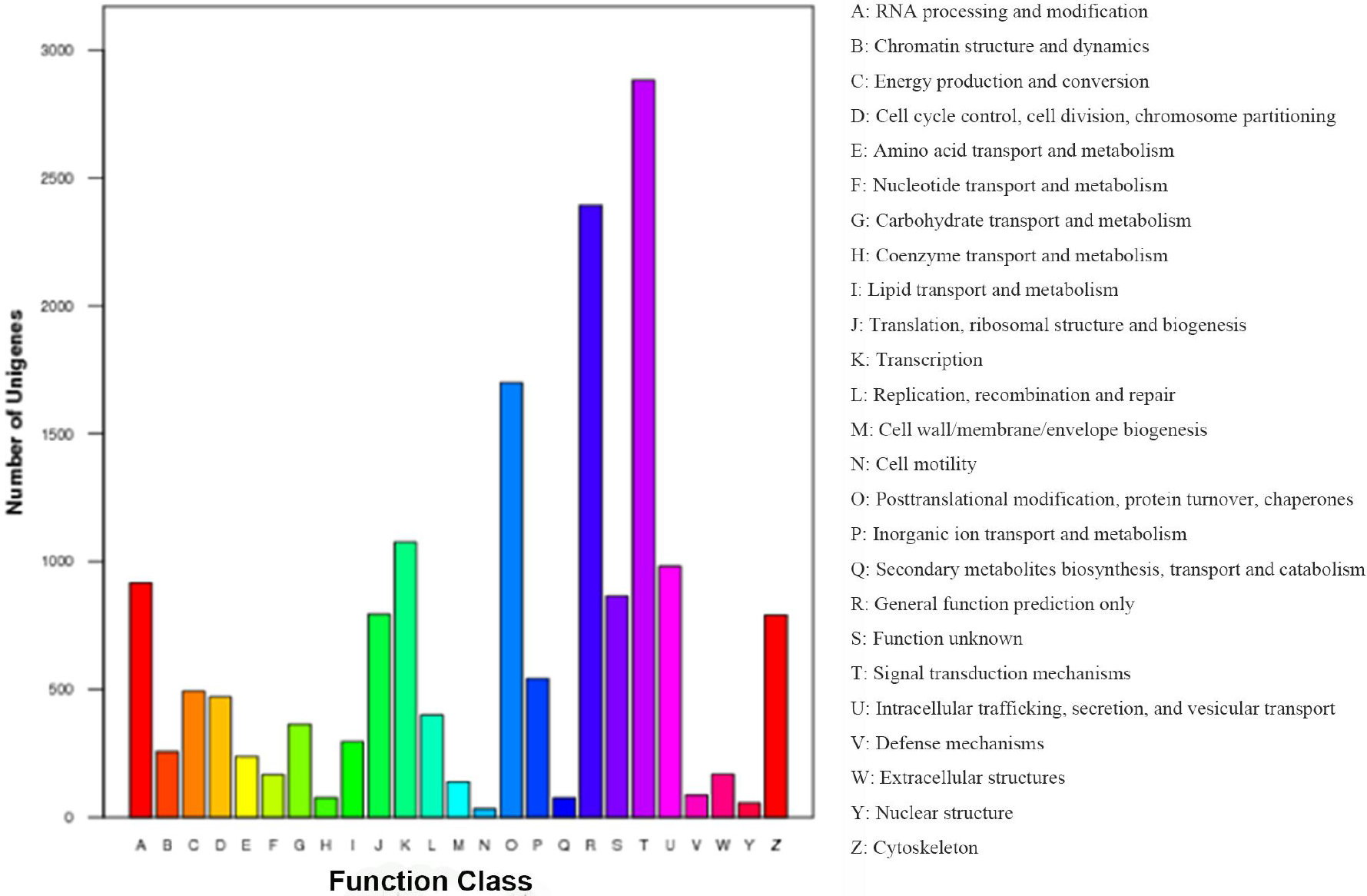
KOG functional annotations of putative proteins among transcripts of *T. pisiformis*

**Figure 4.**
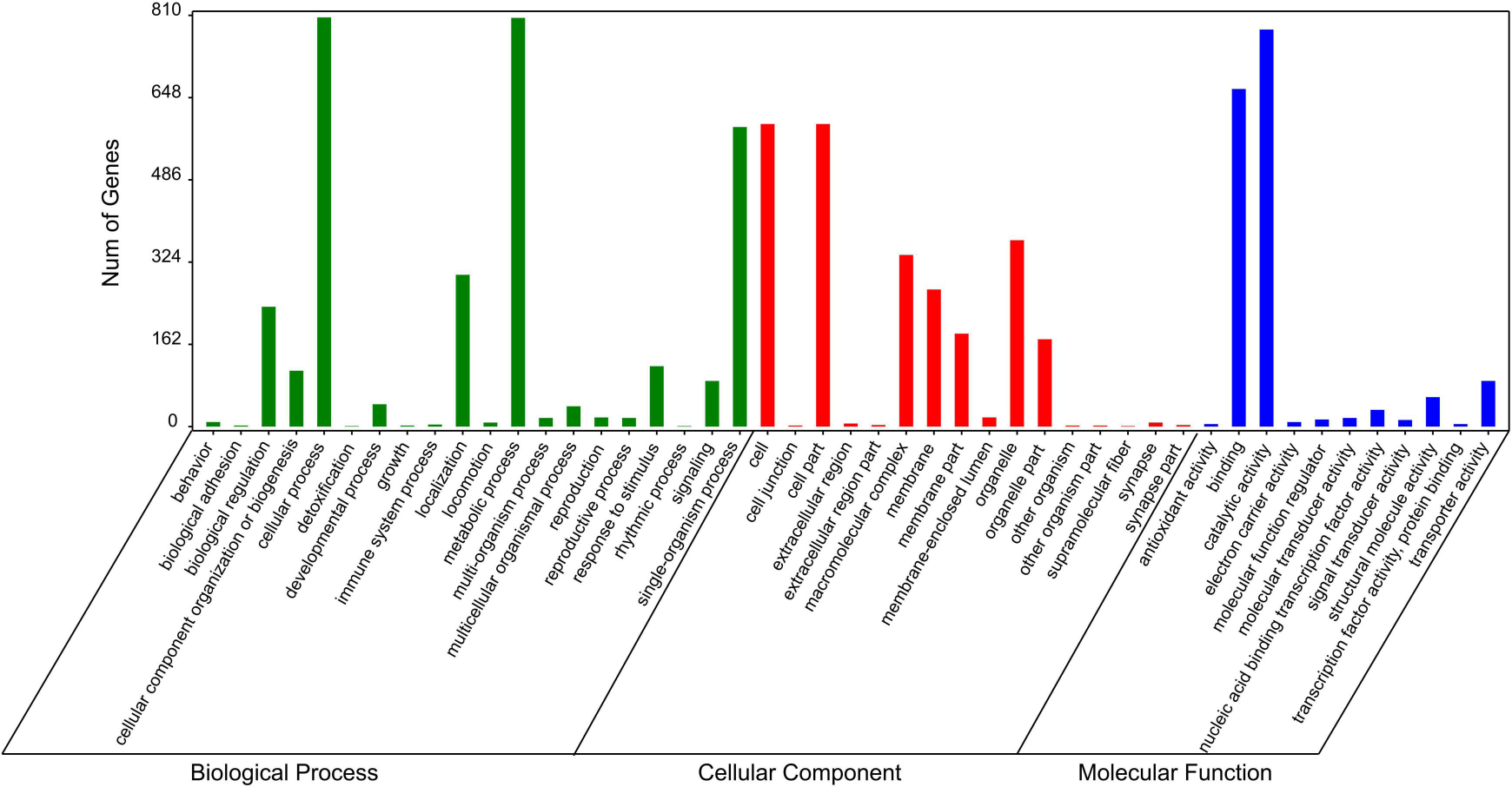
GO annotations of unigenes in *T. pisiformis* transcriptome

### 2.3 Unigene metabolic pathway analysis

A total of 3,122 unigenes were mapped to 227 KEGG pathways. The KEGG pathways included five categories: “metabolism”, “genetic information processing”, “environmental information processing”, “cellular processes” and “organismal systems” (The specific data are not illustrated here). The most abundant pathway was transcription ribosome, including 273 transcriptions, followed by 216 transcriptions in endocytosis and 193 transcriptions in spliceosome pathway (Additional file 2: Table S2). The metabolic pathway and genetic information processing pathway included over 3,500 unigenes. Based on the KEGG pathway, the top 5 KEGG pathways were ribosome (273, 8.74%), endocytosis (216, 6.92%), spliceosome (193, 6.18%), protein processing in endoplasmic reticulum (176, 5.64%) and oxidative phosphorylation (176, 5.64%). These results indicated the active protein metabolism in *T. pisiformis*.

### 2.4 Comparison of gene expression profile of T. pisiformis from different developmental stages

To detect gene expression differences in different development stages in the life cycle of *T. pisiformis*, the researchers analyzed differentially expressed genes in the Cp (larva) groups and Tp (adult) groups. The results suggested that 10,247 DEGs were identified, including 6,910 up-regulated unigenes and 3,337 down-regulated unigenes in the the Tp (adult) compared with Cp (larva) group (Figure 5, Figure 6 and additional file 3: Table S3). DEGs were assigned to 198 KEGG pathways. Among them, “endocytosis” (ko04144, 86 unigenes) and “phagosome” (ko04145, 66) showed significant enrichment (Additional file 4: Table S4). In the pathway enrichment analysis, unigenes were classified into 198 metabolic pathways. The top 10 pathways with the largest unigene numbers were listed in Table 3.

**Figure 5.**
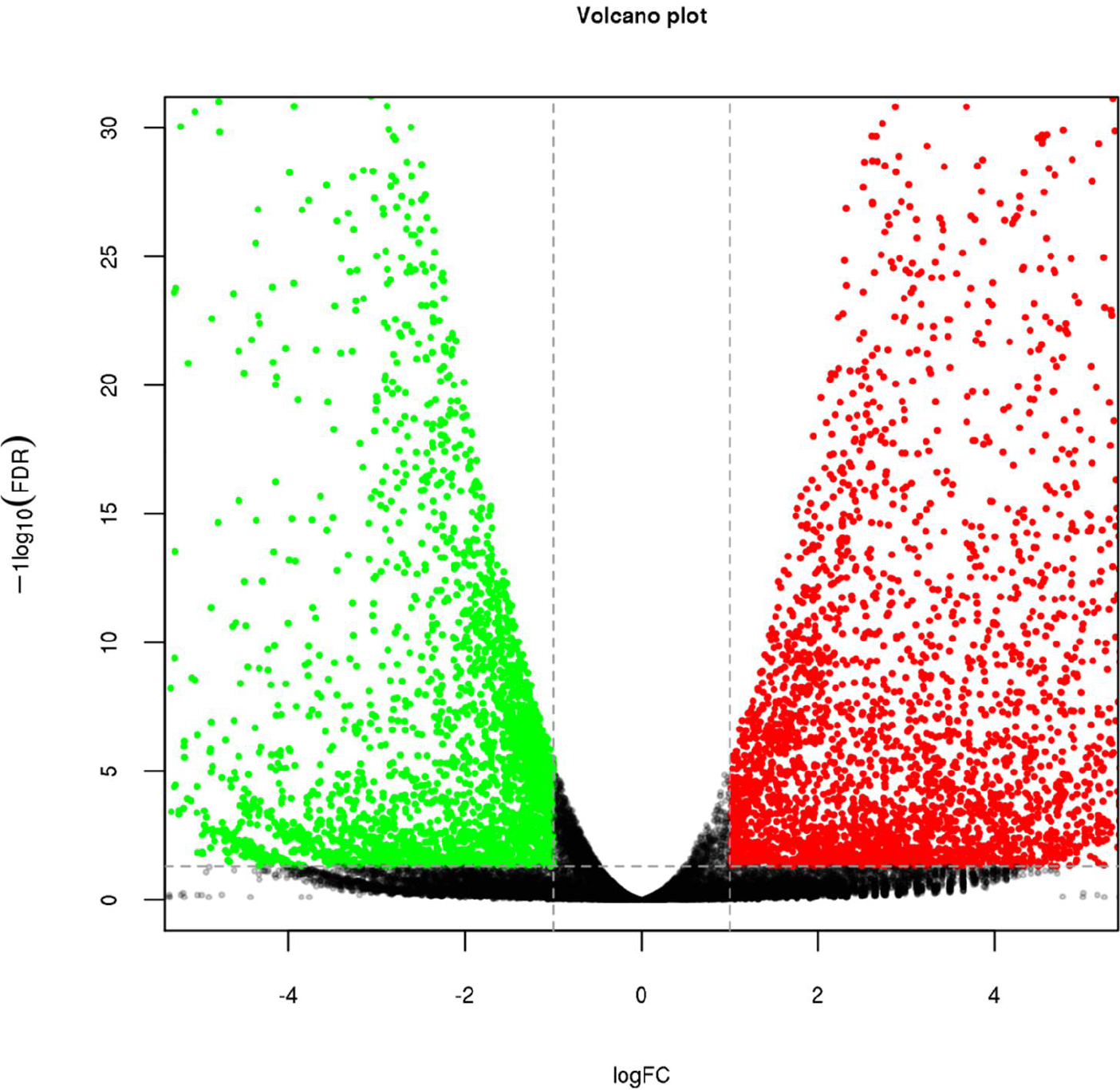
Volcano plot showing differentially expressed genes between three replicates of adult and three replicates of larva of *T. pisiformis*. The x-axis corresponds to the log2 fold-change value, and the y-axis displays the log10 (FDR).The red dots represent the significantly differentially expressed transcripts (p ≤ 0.05 and fold change ≥ 2) between the adult and the larva, while the black dots are not statistically significant (P>0.05).

**Figure 6.**
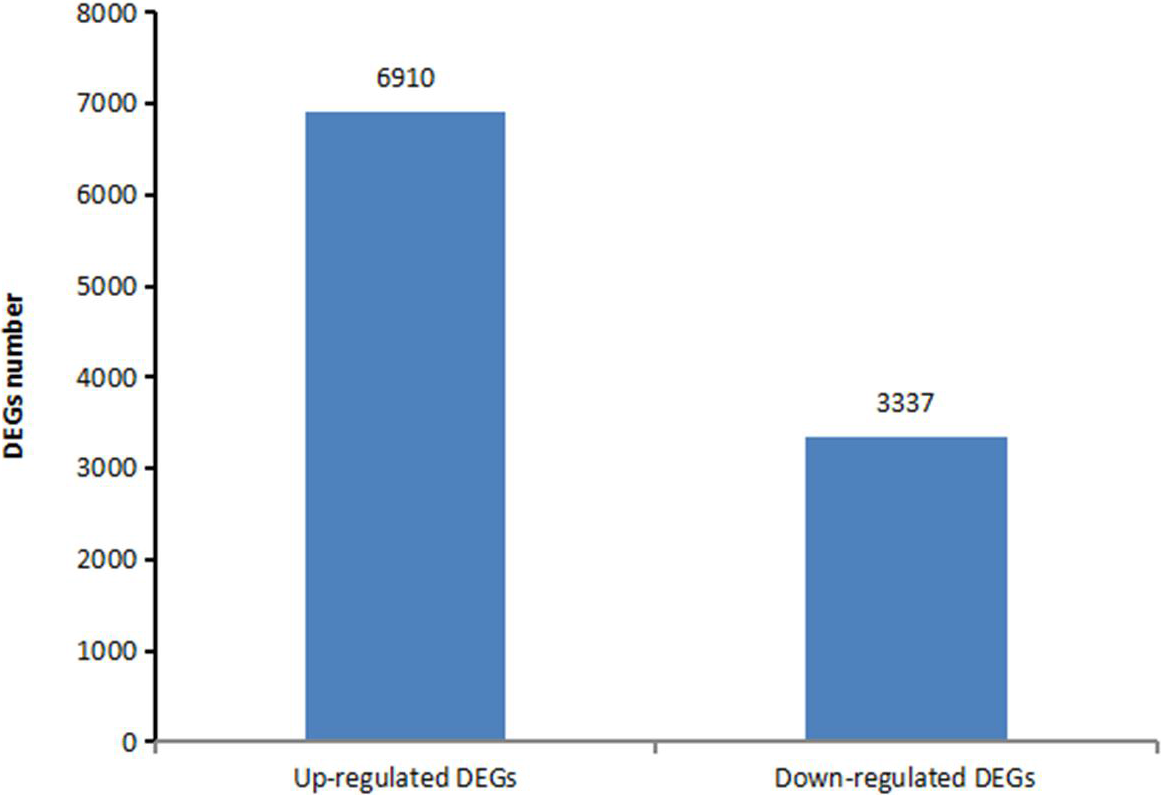
Statistics of differentially expressed genes comparing adult to larva of *T. pisiformis*

**Figure 7.**
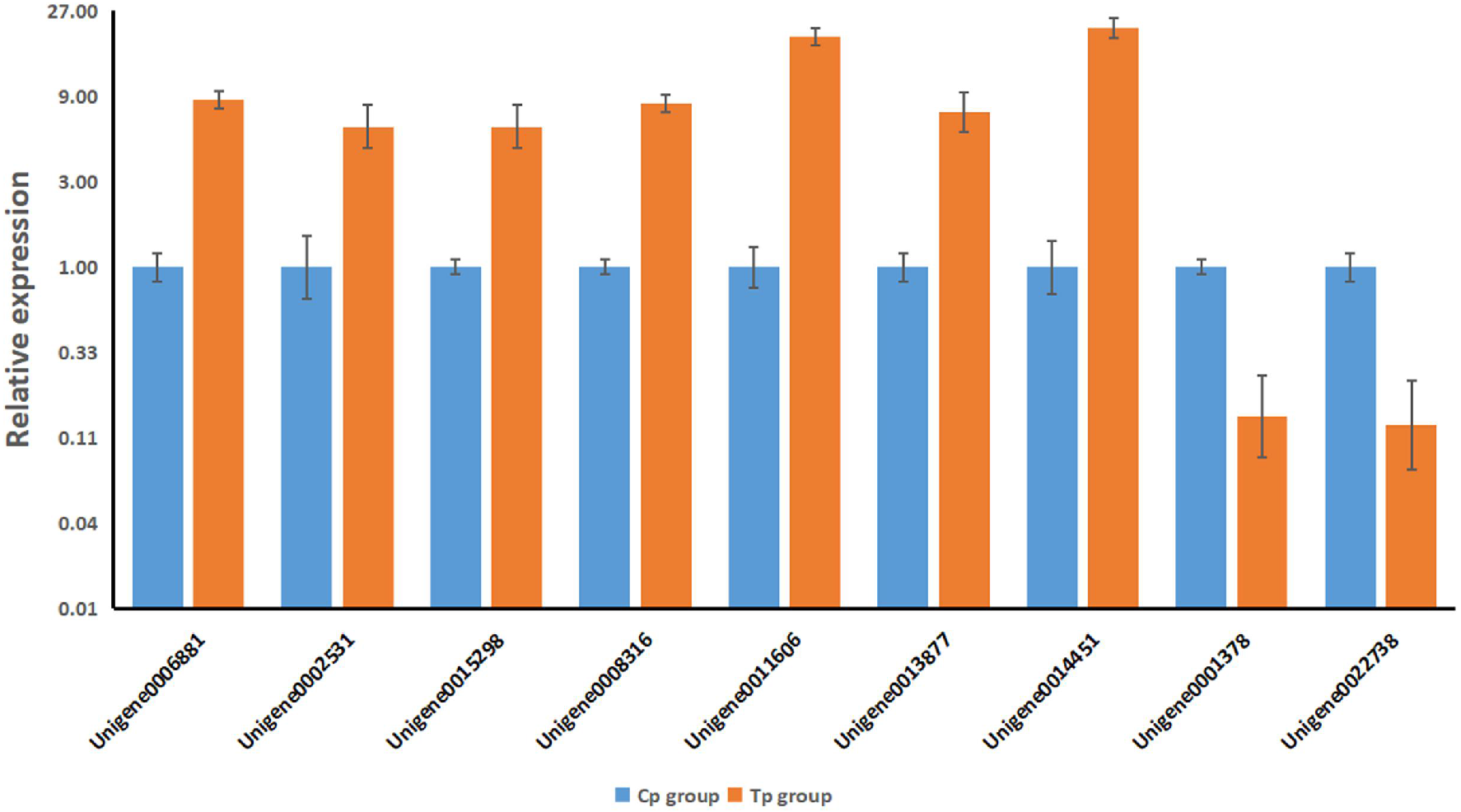
qRT-PCR validation of nine selected genes that were differentially expressed between the Cp group (samples from *T. pisiformis* larva) and the Tp group (samples from *T. pisiformis* adult) in the transcriptomic data. Error bars indicate standard error

**Table 3.**
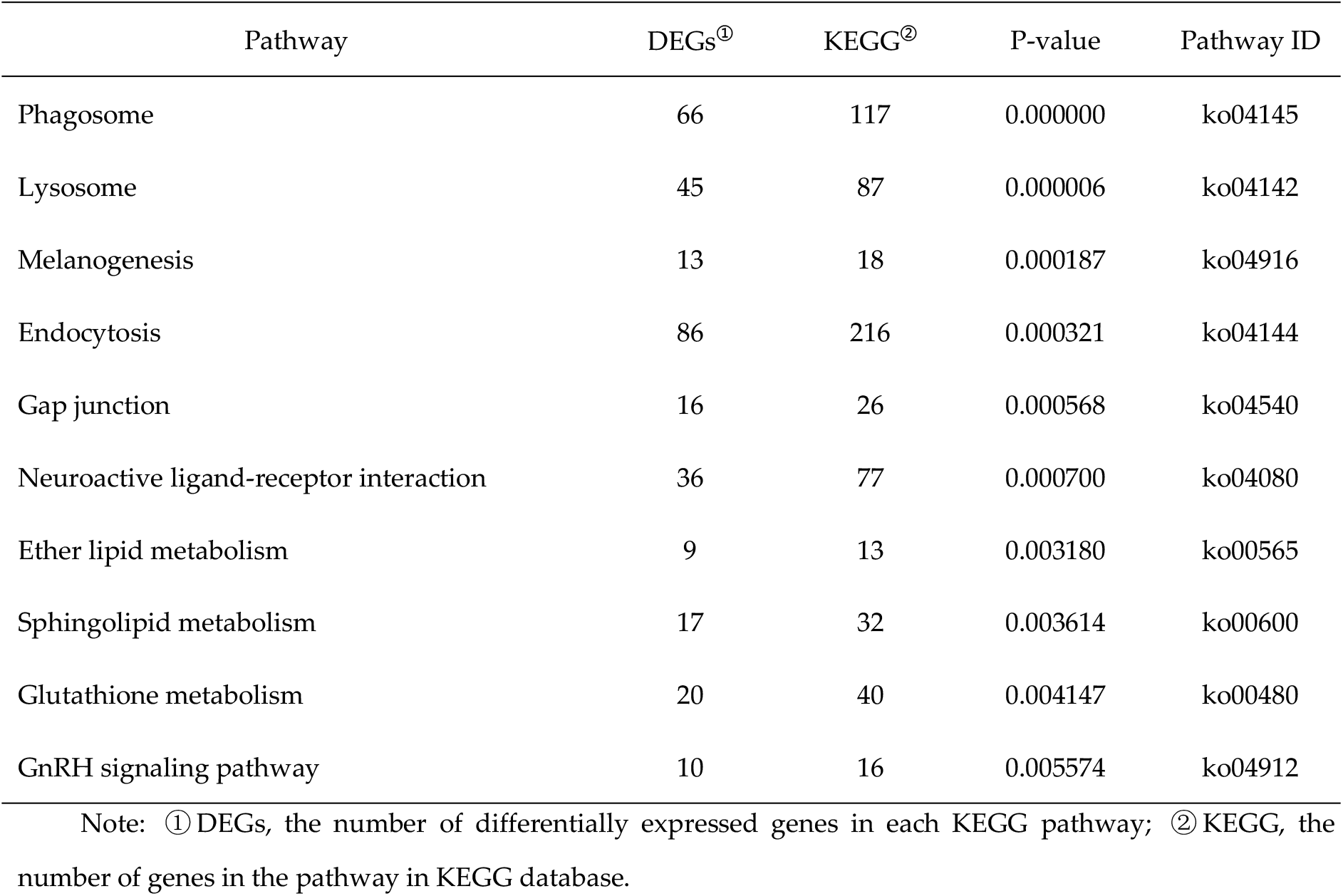
Enriched pathways of differentially expressed genes

| Pathway | DEGs<sup>①</sup> | KEGG<sup>②</sup> | P-value | Pathway ID |
| --- | --- | --- | --- | --- |
| Phagosome | 66 | 117 | 0.000000 | ko04145 |
| Lysosome | 45 | 87 | 0.000006 | ko04142 |
| Melanogenesis | 13 | 18 | 0.000187 | ko04916 |
| Endocytosis | 86 | 216 | 0.000321 | ko04144 |
| Gap junction | 16 | 26 | 0.000568 | ko04540 |
| Neuroactive ligand-receptor interaction | 36 | 77 | 0.000700 | ko04080 |
| Ether lipid metabolism | 9 | 13 | 0.003180 | ko00565 |
| Sphingolipid metabolism | 17 | 32 | 0.003614 | ko00600 |
| Glutathione metabolism | 20 | 40 | 0.004147 | ko00480 |
| GnRH signaling pathway | 10 | 16 | 0.005574 | ko04912 |

Note: ① DEGs, the number of differentially expressed genes in each KEGG pathway; ② KEGG, the number of genes in the pathway in KEGG database.

### 2.5 Development of SSR marker of T. pisiformis

Using microsatellite loci scanning tool MISA, we found that 1,642 SSR loci made up two to six nucleotide repeats in table transcriptome data (Table 4). 1,335 sequences of 36951 contained SSR sites, of which 213 unigenes had more than one SSR loci. Trinucleotide repeat types accounted for the highest proportion, reaching 45.68%, followed by two, four, six and five nucleotide repeat types and the proportions were 35.51%, 13.34%, 2.80% and 2.68%, respectively. In the detection of SSR, a total of 95 kinds of element types were found. The most abundant ten kinds of repeat motifs were AC/GT (375), AGG/CCT (201), ACC/GGT (181), AG/CT (175), AGC/CTG (141), AAG/CTT (67), AAC/GTT (47 A), ATC/ATG (42), AT/AT (32) and ACG/CGT (32). The analysis of the feature of SSR mentioned above is helpful for the development of general molecular markers for *T. pisiformis* and similar species.

**Table 4.** Distribution and compositions of dominant repeat for the different SSR

| Repeat type | SSR number | Proportion (%) | Advantage repeat |
| --- | --- | --- | --- |
| Dinucleotide | 583 | 35.51% | AC/GT |
| Trinucleotide | 750 | 45.68% | AGG/CCT |
| Tetranucleotide | 219 | 13.34% | AGGC/CCTG |
| Pentanucleotide | 44 | 2.68% | ACGAG/CGTCT |
| Hexanucleotide | 46 | 2.80% | ACCGCC/CGGTGG |
| Total | 1642 | 100% |  |

### 2.6 Prediction of allergen genes

The unigenes of *T. pisiformis* in the two development stages were compared by Blastx against allergen protein sequences, and 808 putative allergen genes were identified (Additional file 5: Table S5). There were 618 putative allergen genes producing Blast hits to genes from other *Taenia* and *Echinococcus* (Additional file 6: Table S6), while the other 190 did not produce Blast hits to genes from other *Taenia* and *Echinococcus* (Additional file 7: Table S7).

The upregulated DEGs were similarly analyzed in the adult group relative to the larval group and 343 putative allergen genes were predicted (Additional file 8: Table S8). Among them, 284 were shared with other *Taenia* and *Echinococcus* (Additional file 9: Table S9), and 59 were not (Additional file 10: Table S10). A similar analysis of the significantly downregulated DEGs predicted 461 putative allergen genes (Additional file 11: Table S11), of which 334 produced Blast hits to sequences from *Taenia* and *Echinococcus* (Additional file 12. Table S12), while 127 did not (Additional file 13. Table S13).

### 2.7 Analysising of wnt signaling pathway and wnt genes

To better understand the biochemistry and physiology of *T. pisiformis*, we chose the “wnt signaling” pathway as a case-study for further analysis. Across the transcriptome, this pathway was mapped to 37 unigenes which were grouped into three categories (Canonical pathway, Planar cell polarity pathway and wnt/Ca^2+^ pathway). This pathway was one of the most enriched in downregulated DEGs in the adult group relative to the larvae group; there were 20 downregulated DEGs related to this pathway(Additional file 14. Figure S1). In the results, we found wnt1, wnt4, wnt5a, wnt11 and wnt11b in the larval and adult stage of *T. pisiformis* transcriptome annotations. The wnt2b-a and wnt3a were found in the adult but were absent in the larvae of *T. pisiformis* (Additional file 15. Table S14).

## 3. Discussion

The development of a new generation of high-throughput sequencing technology changed the study of transcriptional studies (Chen and Guo et al., 2017). This is a new method to analyze the complex genomic function and cellular activities of organisms. This method does not need to design probes in advance and can sequence the whole transcriptional activity of any biological growth and development stage under specific conditions. It is more precise and is not subject to cross-hybridization, thereby providing higher accuracy and a larger dynamic range (Wang and Gerstein et al., 2009; 2015). In the current study, the transcriptional sequence of the *T. pisiformis* larvae and adults were sequenced and assembled, using a HiSeq 2000 paired-end sequencing platform and Trinity assembling software. A total of 10,247 DEGs were obtained. The results may lay the foundation for the research about the development of the reproductive system. This results can also provide extensive coverage of the transcriptome in long fragments, which presents large numbers of data to analyze gene expression, predict new genes, and explore metabolic pathways of *T. pisiformis*.

In this study, we carried out transcriptome sequencing analysis of the *T. pisiformis* larvae and adults, and explored the difference of gene expression, metabolic pathway and functional clustering in the two development stages. Sample collection plays an important role in the accuracy and representativeness of the transcriptional data. In order to make the transcript data more comprehensive and representatives, this study carried out three repetitions for each sample to ensure the result.

In 2015, Ju Yan (Ju, 2013) sequenced the mRNA of *E. granulosus* by the Illumina’s Solexa sequencing platform, and obtained 2GB data. In 2017, the transcriptomic of the larva *T. multiceps* was analyzed for the first time. 70,253 unigenes with a mean length of 1492 bp were obtained (Li and Zhang et al., 2017). In 2012, the Illumina sequencing technology was used to detect transcriptome of *T. pisiformis* (Yang and Fu et al., 2012). 72,957 unigenes were assembled. At the same time, the unigenes of *T. pisiformis*, *E. granulosus* and *E. multilocularis* were compared, and the results revealed the distribution characteristics of functional genes. In the current study, reads were assembled into 36,951 unigenes. The assembled unigenes were compared with BLASTX, Nr, Swissprot, KOG and KEGG database. The comparison rates were 34.28%, 22.16%, 20.51% and 17.03%, respectively. 12,844 unigenes were annotated. The average length of *T. pisiformis* unigenes obtained in this study (average length 950 bp) was longer than that in earlier studies (398 bp) (Yang and Fu et al., 2012). This indicates that the sequencing data was well assembled. The unigene annotations included a description of the gene name, analysis of GO terms and metabolic/signaling pathways, which provided biological information at a specific time. Such data could contribute to a more in-depth understanding of gene expression in *T. pisiformis*.

Both diseases and parasites have high correlations with allergy, because of the immunological characteristics that contribute to maintaining the larvae in its host (Vuitton, 2004). Homologous skin-sensitizing antibodies could be detected in the sera of rabbits infected with *T. pisiformis* (Leid and Williams, 1975). In particular, we characterized putative *T. pisiformis* allergen genes and analyzed differences in gene expression between larval and adult stages. This lays the foundation for research about the pathogenic properties of these two stages. Comparisons of *T. pisiformis* DEGs against allergen protein sequences produced 808 predicted allergen genes. Among them, 618 matched sequences in other *Echinococcus* and *Taenia.* Further study on these genes will help researchers understand the interactions between *T. pisiformis* and its hosts.

The wnt signaling pathways are a group of signal transduction pathways involved in a wide range of cellular interactions throughout the development, including regulating cell proliferation, segmentation and axial patterning (Dierick and Bejsovec, 1998). It is known that there are at least three wnt pathways: the canonical pathway, the planar cell polarity (PCP) pathway and the wnt/Ca^2+^ pathway (Logan and Nusse, 2004). Compared with planarians, there are fewer orthologs and paralogs gene of wnt in parasitic flatworms. Riddiford et al. (2011) reported that wnt is more likely to be involved in the evolution of segmentation in platyhelminthes (Riddiford and Olson, 2011). Aulehla et al. (2003) reported that Wnt3a plays a major role in the segmentation clock controlling somitogenesis (Aulehla and Wehrle et al., 2003). Hou et al. (2014) reported that the gene of wnt4 may be related to the process of cysticerci evagination and the scolex/bladder development of *T. solium* cysticerci (Hou and Luo et al., 2014). We found seven wnt genes (wnt1, wnt2b-a, wnt3a, wnt4, wnt5a, wnt11 and wnt11b) in adult stage of *T. pisiformis* transcriptome annotations and five wnt genes in larval stage (Absent of wnt2b-a and wnt3a). The absence of wnt2b-a and wnt3a in larval stage may be related to the somitogenesis. The data of wnt genes in this study may help clarify the role of wnt genes in the development of *T. pisiformis*.

## 4. Materials and Methods

### 4.1 Ethics statement

Animals were handled strictly according to the animal protection laws of the People’s Republic of China (released on Sept. 18^th^, 2009) and the National Standards for Laboratory Animals in China (executed on May 1^st^, 2002). The Animal Ethics Committee of Sichuan Agricultural University (AECSCAU; Approval No. 2014-015) had reviewed and approved this study.

### 4.2 Parasite

*T. pisiformis* larvae were collected from the great omentum of two New Zealand white rabbits naturally infected with this tapeworm from a farm in Sichuan, China. After morphological identification, three larvae were cleaned, labeled and kept in liquid nitrogen, and the other 10 larvae were used for the dog infection. Three adult worms (*T. pisiformis*) were taken out of small intestine, and were washed in warm physiological saline for three times to avoid contamination before they were frozen immediately and stored in liquid nitrogen.

### 4.3 RNA isolation and Illumina sequencing

Total RNA was isolated from single adult (including scoles, neck and strobila) and larva of *T.pisiformis* using Trizol reagent (Invitrogen, Life Technologies, Carlsbad, CA, USA) according to the manufacturer’s protocol. Total RNA of independent samples were stored at −80°C before used. The RNA quality was verified by an Agilent 2100 RNA Nanochip (Agilent, SantaClara, CA, USA) in terms of concentration, RNA integrity number and the 28S:18S ratio.

The OligoTex mRNA mini kit (Qiagen) was used to isolate poly (A) mRNA. Fragmentation buffer was added to interrupt mRNA to shorten fragments (100-400 bp). The first cDNA chain (200±25 bp) was synthesized by six base random primers according to the template of mRNA. Then the second cDNA chains were synthesized by adding buffer dNTPs, RNase H, and DNApolymerasel. Having been purified by QiaQuick PGR kit and eluted with EB buffer, end repair, poly (A) and sequence connection were performed. Then 2% TAE- agarose gel electrophoresis was used to select the size of fragment, and finally PCR amplification. The qualities of the sequencing libraries were assessed on the Agilent Bioanalyzer 2100 system (Agilent Technologies, CA). The library preparations were sequenced on an Illumina HiSeq 2000 platform (Illumina, USA). RNA-Seq data was produced by Guangzhou Jidiao Biotechnology Co. Ltd. For detailed steps, please refer to Wu et al. (Wu and Fu et al., 2012).

### 4.4 Assembly and Annotation

Before assembly, the high-quality clean reads were obtained from raw reads by removing adaptor sequences, duplication sequences, reads containing more than 10% “N‘’ rates (the “N’’ character represents ambiguous bases in reads), and low quality reads containing more than 10% bases with Q-value ≤ 20. All the downstream analyses were performed using clean reads. The Trinity program was used for de novo assembly of the sequence data. These N-free assembly fragments obtained through the overlapping relations among reads are called unigene.

Blastx alignment was carried out between unigenes and databases including NR (NCBI non-redundant protein sequences), KO (KEGG Orthology), SwissProt (a manually annotated and peer-reviewed protein sequence database), GO (Gene Ontology) and COG (the Eukaryotic Ortholog Groups database).

The direction and CDS of unigenes in databases were obtained based on the best alignment results. Unigenes that could not be aligned to the above databases were scanned using the ESTScan software to obtain the CDS and the sequence direction. The MISA program software was applied to analyze the microsatellite loci. Bioinformatics analyses were conducted as previously described (Yang and Fu et al., 2012).

### 4.5 Analysis of gene expression

The Bowtie software was used for analyzing the ratio of comparison. The gene expression amount was estimated by counting the reads numbers mapped to each gene. Expression levels of individual unigenes from different stages in the life cycle of the *T. pisiformis* were evaluated with the method of RPKM (reads per kb per million reads) (He and Xu et al., 2016).

### 4.6 Real-time PCR (qRT-PCR) validation

QRT-PCR was performed to verify the *T.pisiformis* expression data. Primers for differentially expressed genes and the housekeeping gene GAPDH were designed by Primer3 tool and the sequences are available in Additional file 16: Table S15. For qRT- PCR, an ABI7500FAST real-time PCR System (Applied Biosystems, Forster, USA) and a SYBR^®^Premix Ex *Taq*™ II Kit (Takara, Japan) were applied in accordance to manufacturers’ recommendations. The qRT-PCR conditions were 96°C for 2min, followed by 40 cycles of 93°C for 20s and 59°C for 20s. The final melting curve was analyzed. The relative expression level of each gene was calculated using the 2^−ΔΔCt^ method (Livak and Schmittgen, 2001).

### 4.7 Prediction of putative allergens

To predict putative allergens, the unigenes of *T. pisiformis* in each development stage and the differentially expressed unigenes of *T. pisiformis* were compared by Blast against allergen protein sequences from the allergen database website (https://www.uniprot.org).

## Author Contributions

Conceptualization, Lin Chen and Wei Wang; Formal analysis, Lin Chen; Funding acquisition, Wei Wang; Methodology, Lin Chen and Jing Yu; Project administration, Wei Wang; Resources, Lili Ji, Chengzhong Yang and Hua Yu; Supervision, Lin Chen; Writing - review & editing, Jing Xu.

## Acknowledgments

This work was supported by Chengdu University Youth Fund Program [2018XZB06], Sichuan Province Soft Science Program [2014ZR0118] and Sichuan Province Science and Technology Program [2017JY0118].

